# ATP modulates self-perpetuating conformational conversion generating structurally distinct yeast prion amyloids that limit autocatalytic amplification

**DOI:** 10.1101/2022.10.19.512876

**Authors:** Sayanta Mahapatra, Anusha Sarbahi, Neha Punia, Ashish Joshi, Anamika Avni, Anuja Walimbe, Samrat Mukhopadhyay

## Abstract

Prion-like self-perpetuating conformational conversion of proteins into amyloid aggregates is associated with both transmissible neurodegenerative diseases and non-Mendelian inheritance. Here, we demonstrate that ATP modulates the formation and dissolution of amyloids from a yeast prion domain (NM domain of *Saccharomyces cerevisiae* Sup35) and restricts autocatalytic amplification by controlling the amount of fragmentable and seeding-competent aggregates. ATP, at (high) physiological concentrations in the presence of Mg^2+^, kinetically accelerates NM aggregation. Interestingly, ATP also promotes phase-separation-mediated aggregation of a human protein harboring a yeast prion-like domain. We also show that ATP dose independently disaggregates preformed NM fibrils. Furthermore, high concentrations of ATP delimited the number of seeds by generating compact, ATP-bound NM fibrils that exhibited nominal fragmentation by either free ATP or Hsp104 disaggregase. Additionally, (low) pathological ATP concentrations restricted autocatalytic amplification by forming structurally distinct seeding-inefficient amyloids. Our results provide mechanistic underpinnings of concentration-dependent chemical chaperoning by ATP against prion-like transmissions.

## Introduction

Amyloids are proteinaceous, β-sheet-rich ordered assemblies of misfolded proteins that bypass all the surveillance of the protein quality control (PQC) machinery and are often linked with some of the deadly neurodegenerative diseases such as Alzheimer’s, Parkinson’s, prion diseases, amyotrophic lateral sclerosis (ALS), and so on (1)(2)(3). The PQC system is a network of proteins devoted to countering protein misfolding and aggregation, where adenosine triphosphate (ATP) plays an indirect, yet important, role by providing energy to the chaperones involved in protein homeostasis (4). Although only micromolar concentrations of ATP are required for the function of enzymes that include this PQC machinery, the question as to why cells maintain a multifold higher concentration of ATP fascinated researchers to investigate the molecular role of ATP in cells apart from just being the cellular energy currency (5)(6). Intriguingly, in the amyloid deposits isolated from the brain tissues of patients with neurodegenerative diseases, biologically relevant polyanions such as nucleic acids, heparin, and glycosaminoglycans were detected. These reports hinted at a direct interaction between amyloidogenic proteins and the polyanions such as ATP and unveiled a less explored aspect of molecular ATP in protein solubility and aggregation (7)(8)(9).

The amphiphilic ATP molecule consists of the relatively hydrophobic aromatic pyrimidine base connected via a moderately polar ribose sugar unit to a strongly hydrophilic negatively charged triphosphate moiety (Figure 1a). Previous studies showed that ATP alters the aggregation kinetics of various amyloidogenic proteins. For example, it promotes the assembly of acyl phosphatase (AcP), TauK18, amylin, and so on. Whereas, it inhibits the aggregation of eye lens protein γS-Crystallin, synthetic Aβ42 peptide, the prion domain of Mot3, and FUS (Fused in Sarcoma) (10)(11)(12)(5). ATP also disassembles preformed amyloids of FUS at physiologically relevant concentrations (13)(5). Despite several attempts, the chemical nature of interactions of ATP with amyloidogenic proteins or preassembled amyloid aggregates remains poorly understood. In the case of AcP, the electrostatic charge screening by the triphosphate part of ATP was proposed to be responsible for the accelerated aggregation (10). Whereas, for the TauK18 fragment enriched in positively charged residues, the negatively charged triphosphate moiety of ATP promoted fibrillation due to the rapid dimerization via electrostatic crosslinking between lysine residues (11). On the contrary, the solubilization of proteins to inhibit the amyloid formation or the disassembly of existing aggregates is believed to happen due to the hydrotropic properties of ATP by its aromatic moiety, like the typical hydrotropes that are generally used to solubilize hydrophobic compounds (5). However, in another report, this perspective was challenged as ATP was classified as a chaotropic salt in protein solubilization (14). ATP also has shown its ability to alter phase separation, which may control the aggregation of different proteins (15).

**Figure 1.**
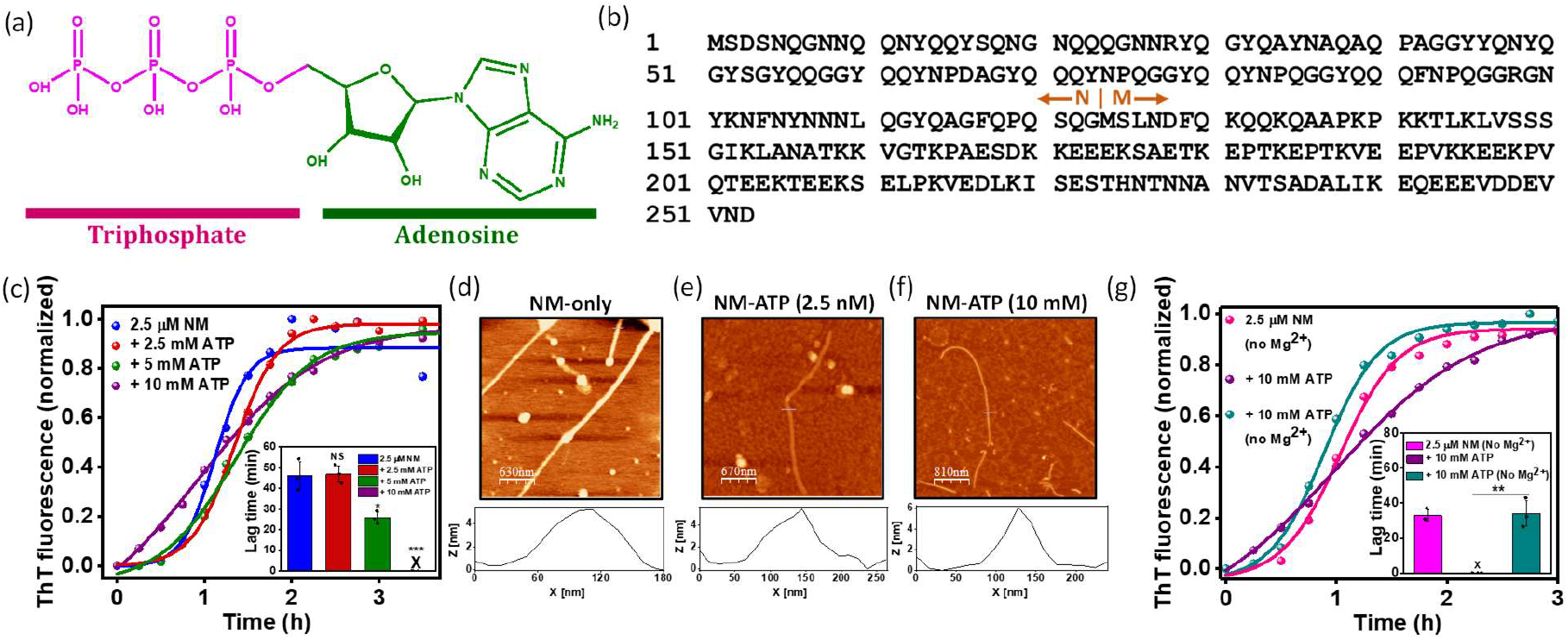
ATP accelerates NM aggregation. (a) The chemical structure of the ATP molecule highlighting the triphosphate (pink) and adenosine (green) moieties. (b) The amino acid sequence of Sup35NM showing the putative boundary between the N-and M-domains. (c) Representative normalized ThT fluorescence kinetics of rotated (80 rpm) NM (2.5 µM) aggregation without or with ATP at room temperature. All reaction mixtures contained 20 mM MgCl_2_ unless otherwise stated. Lag times were retrieved from fitting fluorescence intensities to sigmoidal function. Standard deviations were estimated from three independent replicates (n = 3), NS, *P* ˂ 0.05, *P* ˂ 0.001 for 2.5 mM, 5 mM and 10 mM, respectively, compared to the NM-only aggregation (inset). (d) AFM image of NM-only fibrils (2.5 µM monomers) formed after 6 h of aggregation reaction with a height of ∼6 nm. (e,f) AFM images of ATP-mediated NM fibrils (2.5 µM monomers) aggregated in the presence of (e) 2.5 nM ATP and (f) 10 mM ATP, with a height of ∼ 5 nm. The ATP concentration with which the aggregation of 2.5 µM NM was set up is mentioned in the parenthesis in the case of NM-ATP amyloids. (g) Representative normalized ThT fluorescence kinetics of rotated (80 rpm) NM (2.5 µM monomers) aggregation in the absence of ATP and MgCl_2._ Also, NM (2.5 µM monomer) was polymerized in the presence of 10 mM ATP without or with MgCl_2_ at room temperature and 80 rpm. Lag times were retrieved from three independent replicates (n = 3), *P* < 0.01 for lag time of NM aggregation with ATP and without MgCl_2_ compared to the NM aggregation with MgCl_2_ and ATP (inset). The trace (violet) in Figure 3c (+ 10 mM ATP) is shown in Figure 3g for comparison with and without Mg^2+^.

The unique potential of molecular ATP to control both the formation and dissolution of amyloids might play a pivotal role in regulating prion-like colonization of amyloids, which remains largely uncharacterized. A body of recent evidence suggests that pathological amyloids such as Aβ, tau, α-synuclein, FUS, Huntingtin, p53, and so on follow the prion-like mechanism to invade the uninfected tissues (16)(17)(18)(19)(20). According to this mechanism, preformed amyloid entities known as seeds can be transmitted to the neighboring healthy cells, where they can template amyloid formation (21). Apart from the formation, the fragmentation of high molecular weight fibrils by cellular disaggregases is critical to generate growth-competent amyloids that facilitate prion-like propagation (22)(23)(24)(25). Such fragmented fibrils and oligomers exhibit higher permeability through lipid bilayers of both donor and acceptor cells and are considered as the predominant species for infection and toxicity across amyloid-associated diseases (26)(27)(28). The yeast functional prion protein, Sup35, serves as an excellent model to study the prion-like behavior of transmissible amyloids. The intrinsically disordered prion domain of Sup35 (NM domain) governs the ability of the protein to exist either in soluble or in the amyloid form giving rise to two distinct phenotypes, [*psi*^-^] and [*PSI*^+^], respectively. Furthermore, the cross-generational, non-Mendelian inheritance of [*PSI*^+^] phenotype in budding yeasts can be utilized as the indicator of a prion-like mechanism (29)(30). Additionally, regions comprising positively charged amino acids often linked to prion-like low-complexity domains are associated with physiological functions and diseases. These regions can electrostatically interact with ATP and can be used as archetypes to elucidate its influence in the cascade of seeded amyloid amplification. In this work, we show the ATP-dependent modulation of amyloid aggregation of prion determinant (NM) of a yeast prion protein, Sup35 (Figure 1b). Additionally, phase-separation-mediated aggregation of FUS that contains a yeast prion-like domain exhibited a similar effect in the presence of ATP. We further demonstrate that the ATP controls the prion-like propagation in a concentration-dependent manner by altering the amount and ability of fragmentable and self-templating amyloid seeds.

## Results

### ATP accelerates amyloid aggregation of NM

The aggregation of NM displayed typical nucleation-dependent polymerization kinetics with a lag phase of ∼55 min (31). The lag phase was eliminated in the presence of 10 mM ATP ([NM] = 2.5 µM). However, such an acceleration of aggregation was not observed for the lower concentrations of ATP that showed comparable lag times with respect to the NM-only aggregation (Figure 1c). Next, in order to visualize the nanoscale morphology of these fibrils, we performed atomic force microscopy (AFM). We observed that the NM-only fibrils or NM fibrils formed from ATP-controlled aggregation (NM-ATP fibrils) possess morphological similarities despite their distinct kinetic profiles (Figure 1d-f). Next, we hypothesized that this effect of ATP in accelerating the aggregation is because NM harbors a lot of positively charged amino acid residues, such as lysine. Therefore, these positively charged moieties were possibly electrostatically screened by the polyanion ATP at higher concentrations resulting in speeding up aggregation by reducing the repulsion between precursors. However, our aggregation kinetics studies in the presence of various concentrations of MgCl_2_ revealed that a much higher concentration of MgCl_2_ was required to eliminate the lag phase of NM aggregation compared to the ATP (Figure S1a,b). This corroborated that the charge compensation might not be the reason that altered the fibrillation kinetics. We would like to mention that we used the sodium salt of ATP instead of the magnesium salt and a buffer with 20 mM MgCl_2_ for our studies if not otherwise stated. Over the years, magnesium ions have been reported to be indispensable with ATP as a cofactor to interact with several enzymes (32). Hence, this experimental design permitted us to decipher the role of magnesium ions along with ATP in modulating the aggregation behavior, as we could set up the aggregation in the absence of magnesium ions. Interestingly, without Mg^2+^, ATP did not alter the aggregation of NM, corroborating the importance of the Mg^2+^-ATP complex in promoting the assembly of NM (Figure 1g). These results pointed out that a high amount of ATP, with the help of Mg^2+^, accelerates the NM aggregation that generates fibrils that morphologically resemble the NM-only fibrils.

### ATP promotes phase separation-mediated aggregation of FUS

Next, in order to extend our hypothesis of ATP-mediated acceleration in aggregation, we selected FUS as it harbors a yeast prion-like low-complexity domain as well as a positively charged domain (Figure 2a). Such an amino acid composition makes this protein a potential candidate for electrostatic binding to ATP which can modulate its aggregation. Aberrant aggregation of FUS is implicated in various neurodegenerative diseases, such as amyotrophic lateral sclerosis, frontotemporal dementia, and so on (33)(34)(35). Interestingly, aggregation of FUS is shown to be mediated by liquid-liquid phase separation (36). Thus, to elucidate whether ATP can also affect the aggregation that is mediated by phase separation, we set up phase separation experiments using 10 µM FUS in the presence of a varying concentration of ATP and followed it by turbidity at 350 nm. In the presence of 10 mM ATP, a rapid increase in the turbidity suggested phase separation of FUS. However, in the case of 2.5 mM ATP, there was only a nominal acceleration in the droplet formation. Below that concentration, ATP remained ineffective in changing the time course of droplet formation, as observed in the trend of increase in turbidity with time (Figure 2b). Nevertheless, similar to NM aggregation, ATP was also unable to promote droplet formation in the absence of Mg^2+^, as observed by the turbidity assay (Figure S1c), indicating a complexation between ATP and Mg^2+^ is critical to induce phase separation of FUS. Further, to confirm our turbidity data, we performed confocal microscopy and observed spherical droplets of fluorescently-labeled FUS after 1 min from the commencement of droplet reactions only in the presence of 10 mM ATP along with Mg^2+^. Whereas, In all other instances, the droplets were visible much later (Figure 2c). Next, to probe the maturation of these liquid-like droplets to solid-like aggregates, we monitored ThT fluorescence kinetics. The increase in the ThT fluorescence followed the typical sigmoidal pattern, which corroborated the liquid-to-solid transition of these FUS-ATP droplets that, upon aging, produced aggregates. Thus,10 mM ATP not only accelerated the formation of droplets but also sped up their transition into aggregates, as detected by the reduction in the lag time. This happened possibly due to more liquidity of these droplets as indicated by FRAP recovery that facilitated the oligomerization (15)(37). In the case of 2.5 mM ATP, this reduction was insignificant as the aggregation kinetics almost resembled the FUS-only aggregation (Figure 2d-g, S1d,e). Together, this set of results validated the notion that despite the intricate differences in the aggregation mechanism, similar to NM polymerization, high concentrations of ATP also promoted the aggregation of FUS to generate aggregates via promoting phase separation. Moreover, Mg^2+^, in conjunction with ATP, plays an important role in controlling the phase transition of FUS. Further, a recent report suggested that apart from controlling the aggregation, ATP can also disaggregate preformed fibrils to generate lower molecular weight species (5). As discussed earlier, disaggregation of higher-order amyloids is critical to generate an ample number of seeds for prion-like propagation. Therefore, we next chose NM amyloids over FUS amyloids for our subsequent ATP-induced disaggregation experiments since NM is a more well-studied system to monitor amyloid disaggregation *in vitro* in the context of prion-like mechanism.

**Figure 2.**
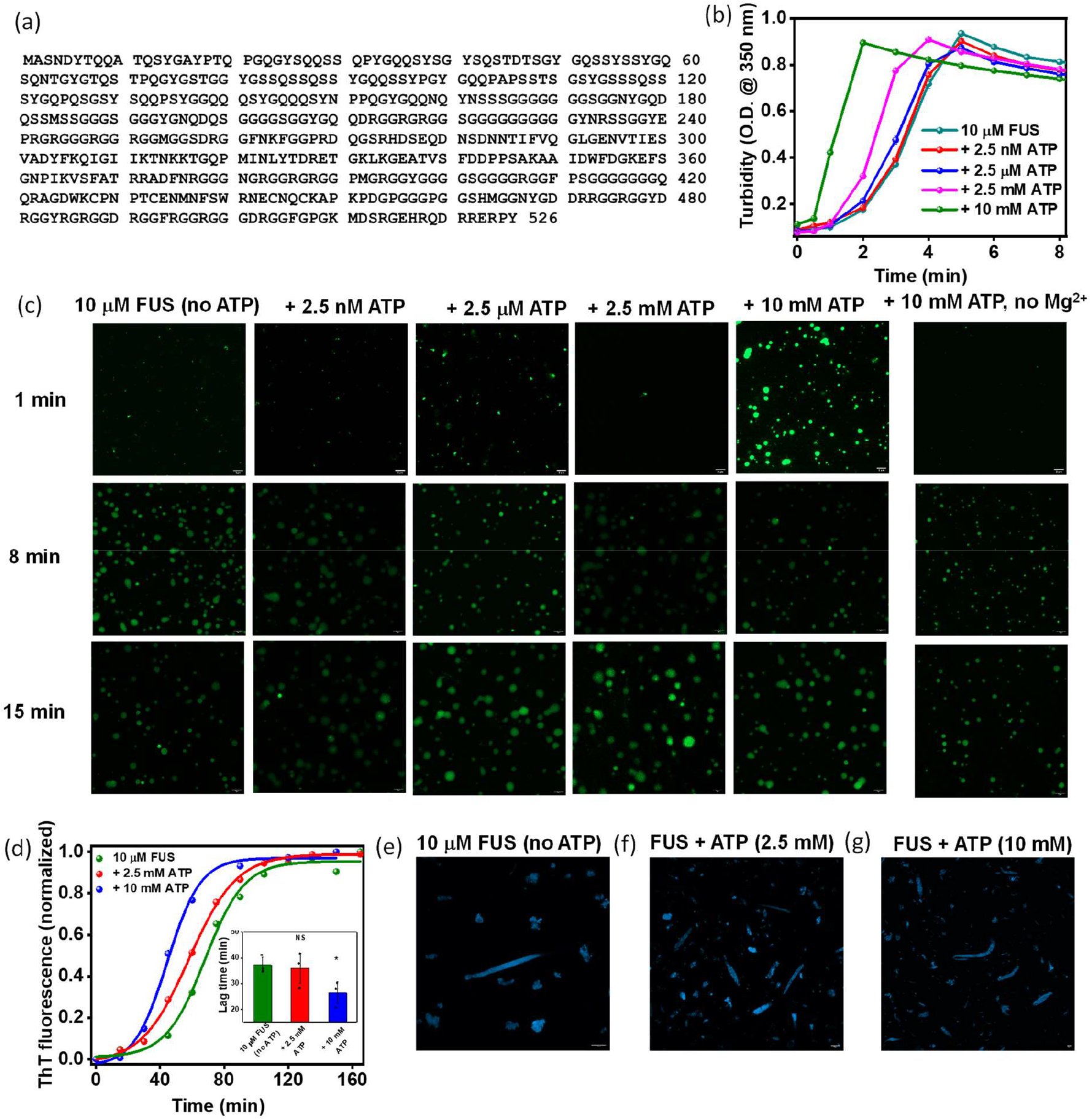
ATP promotes FUS aggregation via phase separation. (a) The sequence of full-length FUS. (b) Liquid-liquid phase separation of FUS in the absence or presence of ATP as monitored by the turbidity assay (at 350 nm). Standard deviations were calculated from three replicates (n = 3). (c) Confocal images of fluorescently labeled FUS droplets (10 µM) at room temperature in the absence or presence of ATP. Confocal images of fluorescently labeled FUS droplets (10 µM) at room temperature in the presence of 10 mM ATP but in the absence of MgCl_2_. (d) Representative normalized ThT fluorescence kinetics of rotated (600 rpm) 10 µM FUS aggregation in the absence or presence of ATP. The reactions were kept in quiescent condition for 15 min to monitor phase separation, as mentioned in Figure 2b,c, and then rotated at 600 rpm for the aggregation. Lag times are shown in the inset. Standard deviations were calculated from three independent replicates (n = 3); NS, *P* < 0.05 for lag times in the presence of 2.5 mM and 10 mM ATP compared to the lag time of the FUS-only aggregation reaction. (e-g) The aggregates formed from rotated (600 rpm) polymerization of (e) 10 µM FUS in the absence of ATP or the presence of (f) 2.5 mM ATP (g) 10 mM ATP as imaged using ThT fluorescence.

### ATP non-stoichiometrically disaggregates preformed amyloids

In order to discern the role of ATP as an amyloid disaggregating agent, we introduced ATP in a wide range of concentrations to the NM-only fibrils in the saturation phase of the aggregation reactions. Interestingly, we observed an immediate drop in ThT fluorescence upon the addition of ATP that indicated the disaggregation of fibrils, which was found to be independent of the presence or absence of MgCl_2_ (Figure 3a and S2c,d). A very low concentration of Hsp104, which can disaggregate amyloids only in higher concentrations, inhibited the ATP-mediated disaggregation probably by hydrolyzing ATP due to its ATPase activity when it was introduced prior to ATP to the fibrils (Figure S2e,f). This observation further confirmed the role of ATP in amyloid dissolution. Next, upon careful analysis of the ThT fluorescence before and after disaggregation, we estimated that the extent of disaggregation remained similar, irrespective of the dose of ATP (Figure 3b). To further establish the non-stoichiometric nature of ATP-amyloid interaction in aggregate solubilization, we sedimented NM amyloids before and after the introduction of ATP. However, we noticed a minimal yet similar decrease in the fraction of sedimentable amyloids after disaggregation, irrespective of the concentration of ATP used (Figure 3c,d). Furthermore, this led us to postulate that there was only a nominal fibril to non-sedimentable amyloid conversion upon the ATP-mediated amyloid disassembly. Therefore, we utilized the AFM imaging in conjunction with immunoreactivity against the A11 and OC antibodies that specifically detects amyloidogenic oligomers and amyloid fibrils, respectively, to precisely characterize the products of the disaggregation caused by ATP (38)(39)(40). Moreover, we compared these particles with the species generated from the disaggregation of NM fibrils by the canonical disaggregating chaperone of yeasts called Hsp104. The AFM images showed that the matured fibrils fragmented into protofibrillar aggregates with reduced length and no spherical particles reminiscent of oligomers by ATP (Figure 3e-h and S2a,b). In contrast, disassembly by Hsp104 resulted in a mixture of oligomers and short protofibrils in AFM (Figure 3i,j). Further, we observed an increase in the A11 signal on Hsp104-induced NM disaggregation, confirming fibril to oligomer formation. As anticipated, the signal for oligomers reduced after the introduction of ATP, validating no generation of oligomers from mature fibrils upon disaggregation (Figure 3k). This was also supported by the immunoblots against OC antibodies that showed no detectable change in signal upon ATP-mediated disaggregation, implying only fibril to shorter fibril conversion (Figure S2g). Taken together, our results showed that ATP can act as a unique, non-canonical disaggregating agent that non-stoichiometrically fragments matured fibrils without yielding oligomers. This was distinct from the case of the disaggregases that generated oligomers that are crucial for prion-like propagation.

**Figure 3.**
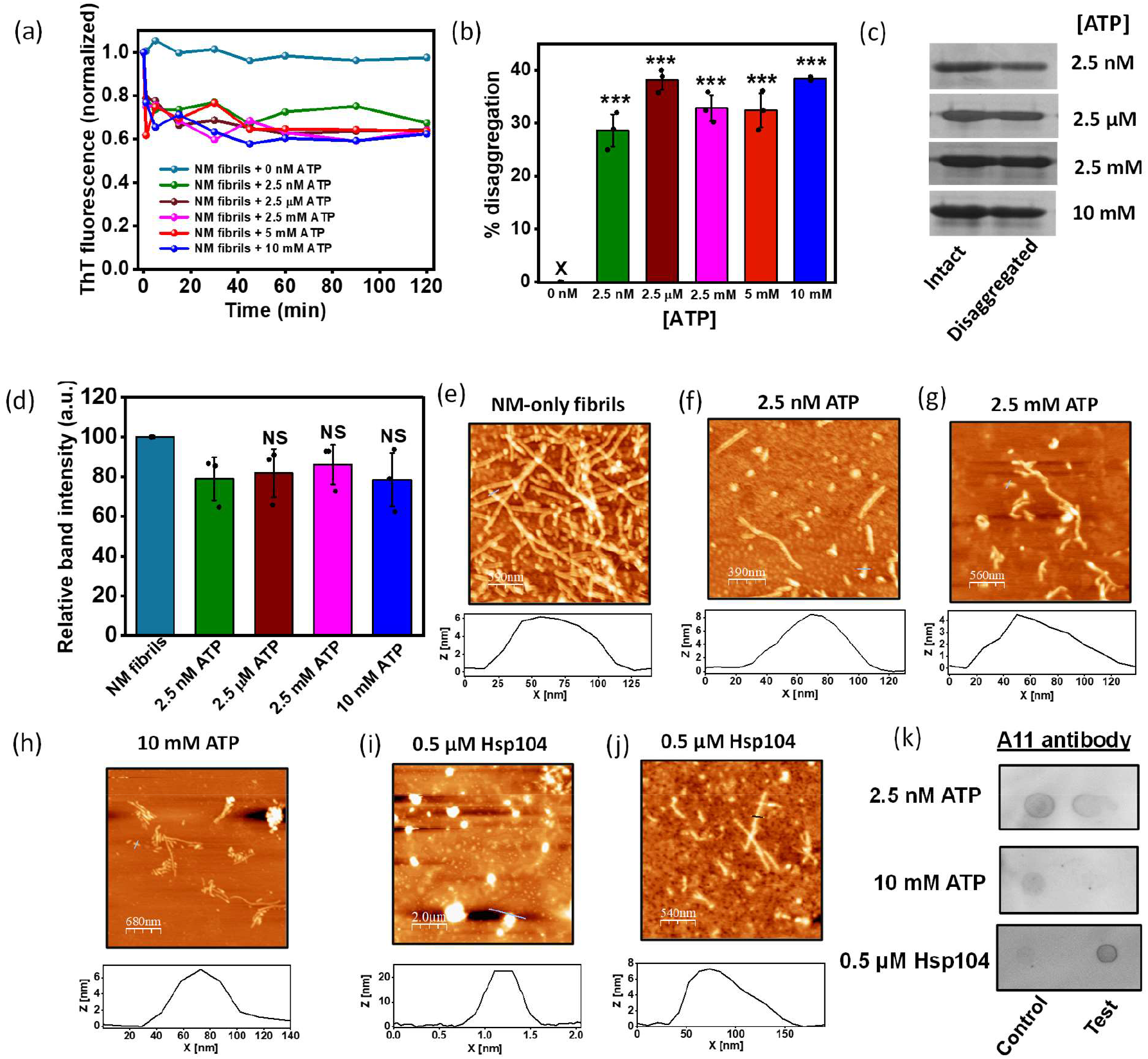
ATP disaggregates matured NM-only fibrils. (a) Representative disaggregation kinetics of NM-only fibrils (2.5 µM monomers) without or with ATP at room temperature and 80 rpm and (b) the percentage of disaggregation. Standard deviations were estimated from three independent replicates (P < 0.001) for the percentage of disaggregation. (c) SDS-PAGE image showing the monomers retrieved before and after disaggregation by ATP of amyloids formed from the rotated (80 rpm) polymerization of NM (2.5 µM) at room temperature after pelleting down amyloids and incubating the pellet in 8 M Urea (20 mM Tris-HCl pH 7.4) overnight. The full uncropped gels are shown in Figure S4. (d) The relative quantification of NM monomers retrieved before and after disaggregation of amyloids by ATP with respect to the band intensity of intact NM-only fibrils in SDS-PAGE by ImageJ software. Standard deviations were calculated from three independent replicates (n = 3); NS, NS, and NS for relative band intensity after disaggregation by 2.5 µM, 2.5 mM, and 10 mM compared to 2.5 nM ATP, respectively. (e) AFM image of intact NM-only fibrils (2.5 µM monomers) formed after 6 h of rotated aggregation (80 rpm) at room temperature with a height of ∼5 nm. (f-h) AFM images of ATP-disaggregated NM amyloids by (f) 2.5 nM ATP (g) 2.5 mM ATP (h) 10 mM ATP with a height of ∼6 nm after 15 min of the introduction of ATP. (i,j) AFM images showing disaggregation of amyloids formed from rotated (80 rpm) aggregation reaction of NM (2.5 µM) at room temperature by Hsp104 (0.5 µM) with the heights of ∼22 nm for oligomers (i) and ∼7 nm for protofibrils (j). (k) Samples from the NM aggregation reactions before and after disaggregation by 2.5 nM ATP, 10 mM ATP, and 0.5 µM Hsp104 were spotted on the nitrocellulose membrane and were probed using the A11 antibody.

### ATP generates compact and stable amyloids

Based on our results described above, we next asked whether ATP can also act as an amyloid disaggregating agent for NM-ATP amyloids similar to the NM-only amyloids assuming their encounter with the free ATP molecules in the cellular milieu. Although all amyloids exhibit a generic cross-β-sheet rich structure, the binding of ATP during aggregation might contribute to the altered packing and kinetic stability of amyloid cores. Hence, assessing the conformational compactness of amyloids is crucial as it may dictate its fragility that drives the prion-like spread by generating amyloid seeds. Accordingly, we intended to characterize the packing of ATP-mediated NM amyloids via its sensitivity against proteolytic digestion by proteinase K (PK). PK is a non-specific endoprotease that digests the constituent monomers of amyloids that are not recruited in the amyloid core. Thus, the digestion pattern of amyloids by PK reveals the supramolecular packing and stability of amyloids (41). Subsequently, we incubated the NM-only or NM-ATP fibrils formed in the presence of different concentrations of ATP with PK and observed that NM-only fibrils were completely digested by it, resulting in lower molecular weight peptides (Figure 4a). On the other hand, NM-ATP amyloids exhibited more resistance to PK digestion showing relatively intense monomeric and sub-monomeric bands in SDS-PAGE. Notably, the NM-ATP fibrils formed in the presence of higher concentrations of ATP exhibited the most resistant amyloid cores demonstrating a prominent, undigested NM monomeric band at ∼35 kDa (Figure 4a). Since NM is rich in positively charged amino acids, we suspected that the electrostatic association between ATP and such residues was crucial for the remarkable kinetic stability of NM-ATP fibrils. To test this hypothesis, the NM-ATP fibrils were treated with the high MgCl_2_ buffer to disrupt the electrostatic interactions between aggregates and ATP prior to the protease digestion. However, in this case, an intermediate protease resistance pattern was observed compared to the NM-only fibrils and the salt-untreated NM-ATP fibrils (Figure 4b). These observations indicated that the electrostatic interactions between the ATP and proteins were only partially responsible for generating stable and compact amyloids. Based on this set of data, we were next interested in deciphering how the stability of NM-ATP fibrils impacted their fragility. Accordingly, we used two different amyloid disassembling agents: (i) the canonical disaggregase enzyme Hsp104 which is known to fragment amyloid fibrils and (ii) the small-molecule amyloid-solubilizer, ATP that was established as an amyloid disaggregating agent for NM-only amyloids by us in the previous section. Intriguingly, the NM-ATP fibrils formed in the presence of higher ATP concentrations showed much less fragmentation propensity, regardless of the disaggregating agent. However, the NM-ATP fibrils created in the presence of a trace amount of ATP demonstrated fragility similar to the NM-only fibrils (Figure 4c-g). In summary, these results indicated that the presence of high concentrations of ATP during the assembly of NM might give rise to stable and compact amyloids that displayed minimal fragmentation and may restrict the prion-like propagation of amyloids by limiting the formation of lower-order seeds.

**Figure 4.**
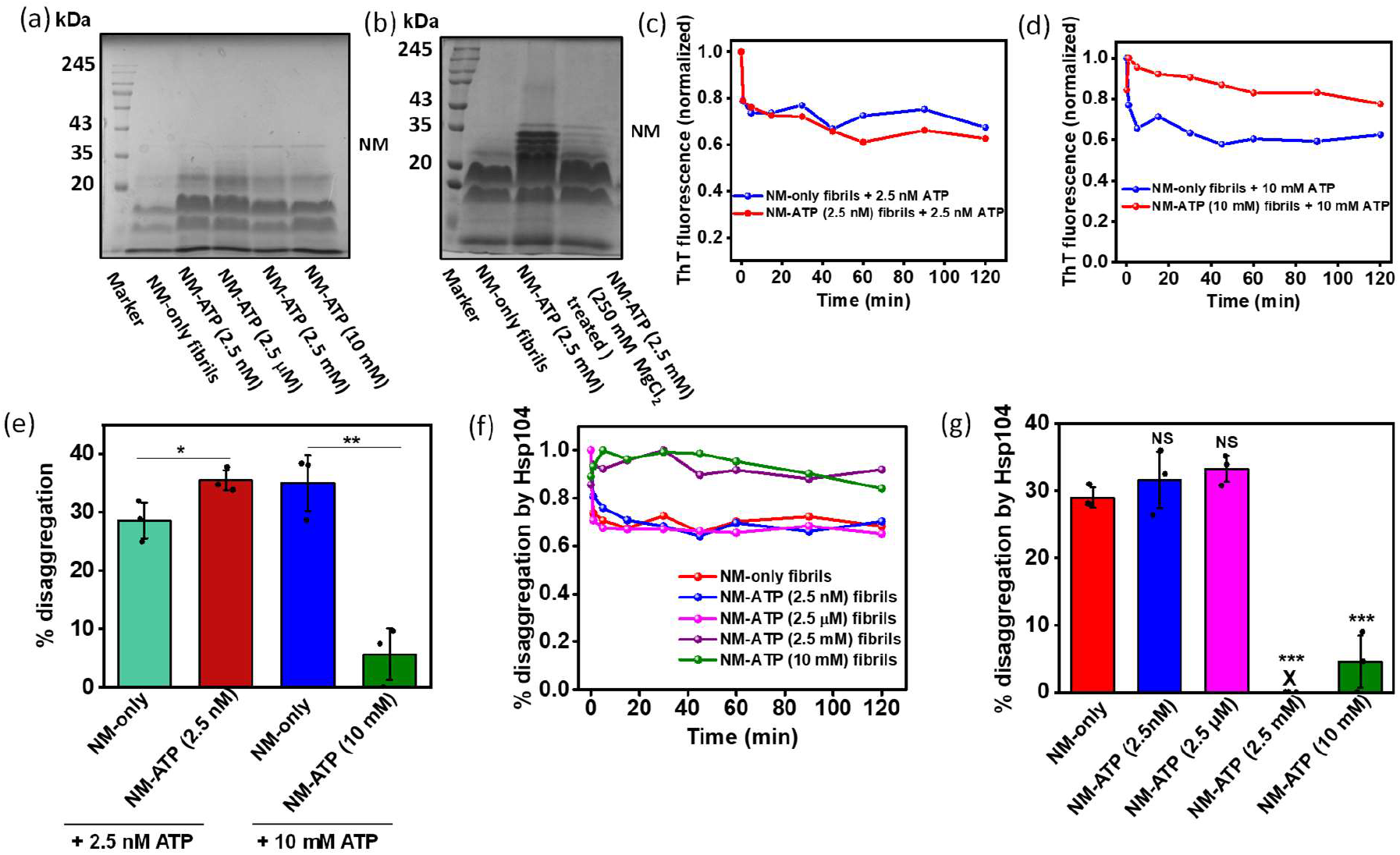
The stability and fragility of NM-ATP amyloids. (a) The concentrated fibrils formed from monomeric NM (2.5 µM) without or with ATP were incubated at 37 ºC for 30 min with proteinase K (PK) (NM: PK 1000:1), followed by SDS-PAGE analysis. The full uncropped gel is shown in Figure S4. (b) The concentrated fibrils formed from monomeric NM (2.5 µM) in the absence or presence of 2.5 mM ATP were incubated at 37 ºC for 30 min with PK (NM: PK 1000:1). Also, the NM-ATP fibrils formed in the presence of 2.5 mM ATP were treated with 250 mM MgCl_2_ and then incubated at 37 ºC for 30 min with PK (NM: PK 1000:1) followed by SDS-PAGE analysis. The full uncropped gel is shown in Figure S4. (c) Representative ThT fluorescence disaggregation kinetics of NM-only fibrils (2.5 µM monomers) or NM-ATP (2.5 nM) fibrils by 2.5 nM ATP at room temperature and 80 rpm. (d) Representative ThT fluorescence disaggregation kinetics of NM-only fibrils (2.5 µM monomers) or NM-ATP (10 mM) fibrils by 10 mM ATP at room temperature and 80 rpm. (e) Percentage disaggregation of NM-only fibrils and NM-ATP (2.5 nM) fibrils by 2.5 nM ATP or NM-only fibrils and NM-ATP (10 mM) fibrils by 10 mM ATP. Standard deviations were calculated from three independent replicates (n = 3), *P* < 0.05, *P* < 0.01 for percentage disaggregation by 2.5 nM ATP of NM-ATP (2.5 nM) fibrils compared to NM-only fibrils and by 10 mM ATP of NM-ATP (10 mM) fibrils compared to NM-only fibrils, respectively. (f) Representative ThT fluorescence disaggregation kinetics of NM-only fibrils (2.5 µM monomers) or NM-ATP fibrils by Hsp104 (0.5 µM), ATP (5 mM), and ATP regeneration system and (g) the percentage of disaggregation. Standard deviations were calculated from three independent replicates (n = 3), NS, NS, *P* < 0.001, *P* < 0.001 percentage disaggregation by Hsp104 of NM-ATP fibrils formed in the presence of 2.5 nM, 2.5 µM, 2.5 mM, and 10 mM ATP compared to NM-only fibrils, respectively.

### ATP at low concentrations creates seeding-inefficient amyloids

Apart from the fragility of amyloids that regulate the number of seeds, the seeding efficiency accessed by their ability to accelerate a fresh aggregation reaction is a critical factor in prion-like amyloid transmission. Hence, we introduced them in the fresh aggregation reactions to elucidate the seeding potential of NM particles aggregated in the presence of ATP. The NM-ATP fibrils polymerized in the presence of millimolar concentrations of ATP exhibited seeding capability similar to the NM-only fibrils. On the contrary, the fibrils aggregated in the presence of low concentrations of ATP failed to accelerate the fresh aggregation reactions, as evident by similar lag times of unseeded and seeded aggregation kinetics (Figure 5a,b). We then intended to reverse the seeding inefficiency of NM-ATP fibrils generated in the presence of very low concentrations of ATP by treating them with a high MgCl_2_ buffer to dissociate the electrostatically attached ATP molecules to amyloids. However, we were only able to partially recover the seeding capability with respect to the NM-only fibrillar seeds validating that amyloid-ATP interaction was not the sole factor that controlled the seeding inability of NM-ATP amyloids (Figure 5c,d). Next, we hypothesized that the fibrils disaggregated by ATP might also efficiently seed the fresh aggregation reaction due to the increase in the number of growth-competent amyloid ends and might facilitate the prion-like propagation. Therefore, we performed seeded aggregation with ATP-disaggregated NM fibrils, and, as the control, the seeds obtained from the disaggregation of NM fibrils by ultrasonic sound pulses. On careful inspection of the seeded aggregation kinetics, we found that the amyloids derived upon disaggregation of fibrils by ATP displayed a seeding ability similar to the intact fibrils. This data revalidated our previous observations in AFM and immunoblots that suggested a minimal change in the number and molecular weight of amyloids upon disaggregation by ATP. In contrast, a much greater seeding ability for sonicated fibrils was observed, indicating a considerable increase in the number of particles upon sonication that eliminated the lag time in seeded aggregation (Figure 5e,f). Taken together, these data indicated that in the millimolar concentration regime, the presence of ATP during aggregation hardly impacted the seeding potential of the NM-ATP fibrils. On the contrary, the trace amount of ATP may lead to the formation of seeding-inefficient amyloids, which can impair their amplification cascade. In addition, unlike other disaggregating agents, the disaggregation of fibrils by ATP did not translate into an improved seeding, unmasking a propagation-neutral attribute of molecular ATP.

**Figure 5.**
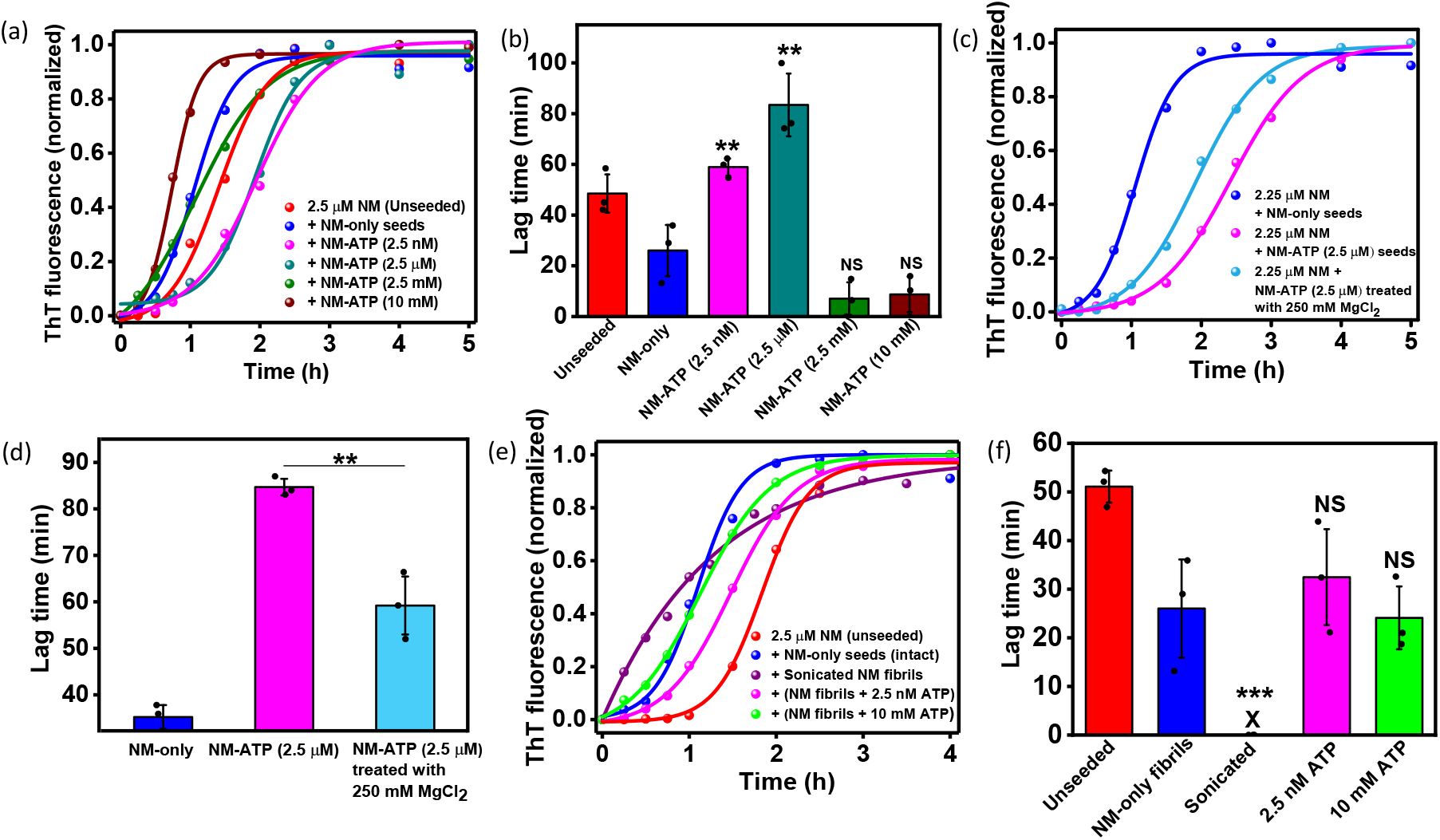
Seeding by NM-ATP amyloids. (a) Representative normalized ThT fluorescence kinetics of rotated (80 rpm) NM aggregation seeded without or with 10 % (w/w) NM-only fibrils (2.5 µM monomers) or NM-ATP fibrils aggregated in the presence of ATP. (b) Lag times estimated from the fitting. Standard deviations were calculated from three independent replicates (n = 3), *P* < 0.01, *P* < 0.01, NS, and NS for lag times in the case of NM-ATP fibrils formed in the presence of 2.5 nM, 2.5 µM, 2.5 mM, and 10 mM ATP compared to NM-only fibrils. (c) Representative normalized ThT fluorescence kinetics of NM aggregation rotated at 80 rpm at room temperature seeded with 10 % (w/w) NM-only fibrils (2.5 µM monomers) or NM-ATP (2.5 µM) fibrils. Also, NM-ATP (2.5 µM) fibrils treated with 250 mM MgCl_2_ were used as seeds. (d) Lag times for seeded aggregation reactions, as mentioned in Figure 5c. Standard deviations were calculated from three independent replicates (n = 3), *P* < 0.01 for the lag time of seeded aggregation with 250 mM MgCl_2_-treated NM-ATP fibrils compared to 250 mM MgCl_2_-untreated NM-ATP fibrils. (e) Representative normalized ThT fluorescence kinetics of NM aggregation reaction rotated at 80 rpm and room temperature in the absence or presence of 10 % (w/w) seeds of intact NM-only fibrils (2.5 µM monomers), sonicated NM fibrils, NM-only fibrils disaggregated by 2.5 nM and 10 mM ATP. (f) Lag times estimated from the fitting. Standard deviations were calculated from three independent replicates (n = 3), *P* < 0.001, NS, NS for seeded aggregation reactions seeded by sonicated NM fibrils, disaggregated NM-only fibrils by 2.5 nM ATP, and 10 mM ATP, respectively, compared to intact NM-only fibrillar seeds. The trace (blue) in 5a, 5c, and 5e are from the same dataset (10 % intact NM-only seeds; total [NM] = 2.5 µM) shown for comparison.

### Conformational characterizations of ATP-mediated amyloids

The presence of molecular ATP during the polymerization of NM altered the stability and seeding potential of the amyloids. However, our PK sensitivity assay or seeded aggregation kinetics of NM-ATP amyloids indicated that the dissociation of electrostatically attached ATP could not completely reverse the consequences of ATP binding. These observations lead to the assumption that some changes may have occurred in the amyloid structure upon binding of ATP to the building blocks of these aggregates, which governed their conformational stability and seeding ability. To capture these plausible conformational remodeling of NM amyloids in the presence of various ATP concentrations, we employed vibrational Raman spectroscopy, which is widely used to gain structural insights into several amyloids (42) (43) (44). We first validated the binding of ATP with amyloids by pelleting down NM-ATP fibrils and recorded Raman spectra (Figure 6a). The prominent dose-dependent peaks of adenine and phosphate moieties at 725 cm^−1^ and 1127 cm^−1^, respectively, of ATP, indicated its binding with fibrils, which otherwise should not sediment with high molecular weight aggregates (Figure 6b,c) (45). In continuation, we also tried to decipher the role of Mg^2+^ in the ATP-amyloid interaction, as our aggregation kinetics data indicated that without Mg^2+^, ATP was unable to exert any effect on NM aggregation. Accordingly, we observed a relatively weak signal of adenine and phosphate moieties for the NM-ATP aggregates formed in the absence of Mg^2+^ compared to the signal obtained from the NM-ATP amyloids generated with Mg^2+^ (Figure 6d-f). This suggested a decreased ATP binding with amyloids in the absence of Mg^2+^. Next, we focused on the amide Ⅲ and amide I regions of Raman spectra, as the positions and widths of these bands between 1230–1320 cm^−1^ and 1630–1700 cm^−1^, respectively, are influenced by the backbone conformations of the polypeptide chains. However, to capture the conformational characteristics of NM-ATP amyloids, we relied mainly on the amide I region, as the amide III region overlapped with the peaks contributed by ATP. The amide I band, which mainly arises due to >C=O stretching of amide, is sensitive to the structural differences in amyloids and reports the secondary structural elements of amyloids formed in the presence of different ATP concentrations. In the case of 10 mM ATP, there was a narrowing of the amide I band that indicated a more ordered, β-sheet-rich secondary structural content in the NM-ATP amyloids compared to the NM-only amyloids corroborating our protease digestion data. However, this narrowing was absent in the case of NM-ATP fibrils aggregated in the absence of Mg^2+^, indicating only a minor structural remodeling by ATP without Mg^2+^, possibly due to reduced ATP binding with amyloids. On the other hand, in the presence of 2.5 µM and 2.5 mM ATP during aggregation, there was a broadening of the amide I band that revealed a structural heterogeneity having a greater proportion of disordered conformation for these amyloids compared to the NM-only amyloids. Additionally, the Raman shift and change in the intensities at 1607 cm^−1^ compared to the NM-only amyloids pointed to the interaction with the tyrosine or lysine of NM with ATP during the aggregation that was again not detected in the absence of Mg^2+^ (Figure 6g,h) (46). We also performed circular dichroism (CD) spectroscopy to reconfirm the secondary structural content of NM-ATP amyloids. We noticed a drop in ratiometric mean residue ellipticity (θ_218_/θ_200_), representing the propensity to form β-sheet at the expense of random coil in the case of NM-ATP fibrils corresponding to nanomolar and micromolar ATP (47). Contrastingly, in the presence of 10 mM ATP, θ_218_/θ_200_ was more for NM-ATP fibrils compared to the NM-only fibrils indicating a greater extent of β-sheet structure in these fibrils corroborating with our Raman data (Figure 6i). We also observed an increase in the fluorescence steady-state anisotropy with increasing ATP concentrations indicating more restricted rotation of the fluorophore in the M-domain due to the more ordered conformation validating our Raman, CD, and proteinase K sensitivity assay results (Figure S3). Taken together, our Raman, CD, and fluorescence anisotropy results provided evidence of the Mg^2+^-dependent conformational remodeling of NM amyloids by ATP, giving rise to altered secondary structures that can potentially govern several biologically important aspects.

**Figure 6.**
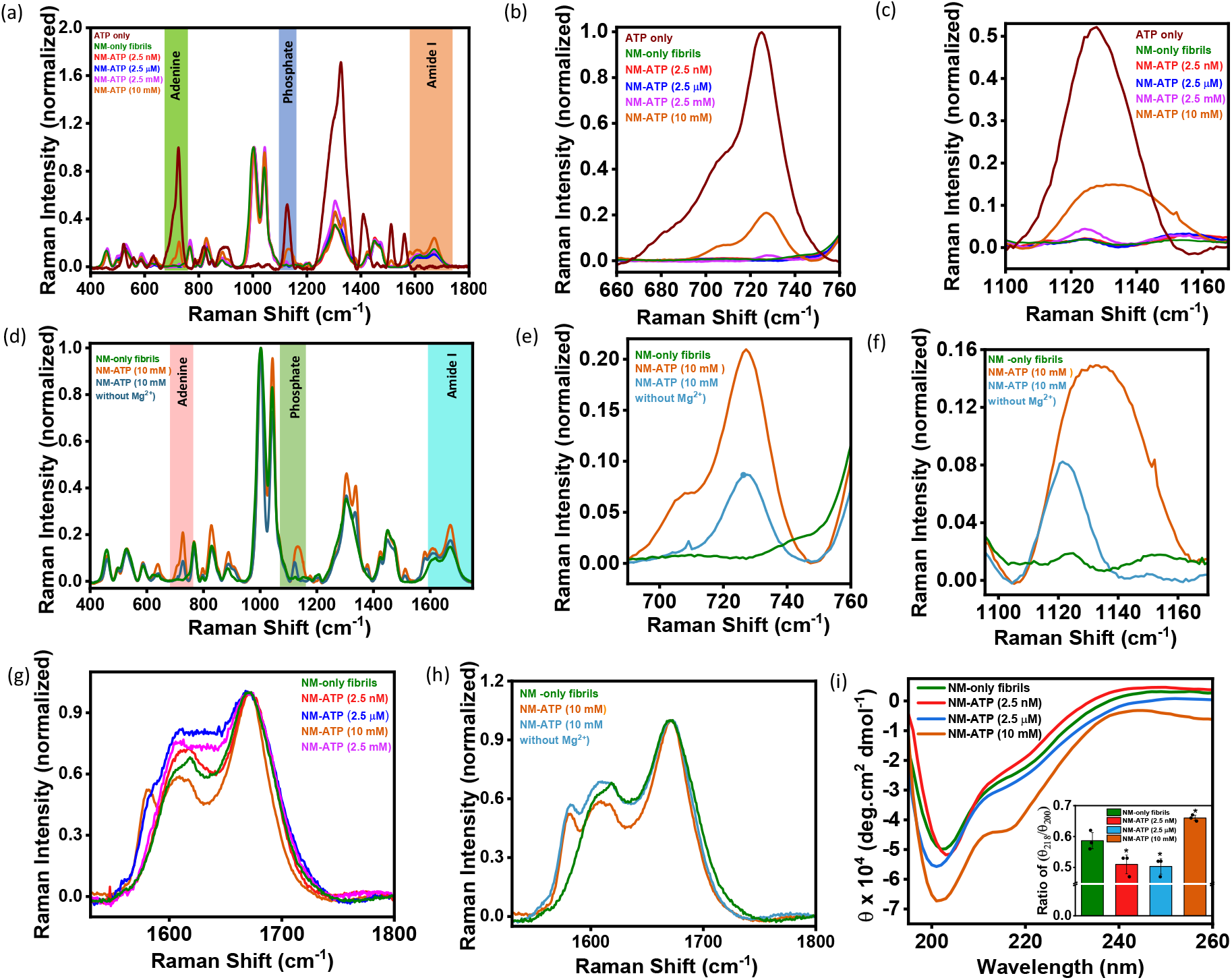
Conformational fingerprinting of NM-ATP amyloids. (a) Raman spectra of NM-only fibrils (2.5 µM monomers) and NM-ATP fibrils formed in the presence of ATP with 2.5 µM NM monomers highlighting the adenine (green), phosphate (blue), and amide I (brown). (b,c) The peaks of adenine and phosphate moieties are shown at (b) 750 cm^−1^ and (c) 1120 cm^−1^, respectively, for NM or NM-ATP fibrils. (d) Raman spectra of NM-only fibrils (2.5 µM monomers) or NM-ATP (10 mM) fibrils in the absence or presence of 20 mM MgCl_2_ highlighting the adenine (pink), phosphate (olive green), and amide I (cyan). (e,f) The peaks of adenine and phosphate moieties are shown at (e) 750 cm^− 1^ and (f) 1120 cm^−1^, respectively, for NM-only fibrils or NM-ATP fibrils formed in the absence or presence of MgCl_2_ (Figure 6a-f are normalized with respect to the phenylalanine at 1002 cm^−1^). (g) The amide 1 region of NM-only or NM-ATP fibrils, as highlighted in brown in Figure 6a. (h) The amide I region of NM-only or NM-ATP fibrils, as highlighted in cyan in Figure 6d. (i) Representative Far-UV CD spectra of NM-only fibrils (2.5 µM monomers) and NM-ATP fibrils in the presence of different concentrations of ATP. The inset shows a plot of ratiometric ellipticity (θ_218_/θ_200_) for NM-only and NM-ATP fibrils. Standard deviations were calculated from three independent replicates (n = 3); *P* < 0.05, *P* < 0.05, and *P* < 0.05 for NM-ATP amyloids formed in the presence of 2.5 nM, 2.5 µM and 10 mM ATP, respectively, as compared to NM-only fibrils (Figure 6g-i are normalized with respect to the amide I intensities).

## Discussion

ATP is well-known for its role as cellular energy currency to fuel various physiological processes. In this current study, we were able to delineate a unique role of ATP in amyloid formation and disaggregation that may regulate the prion-like transmission of self-replicable amyloid entities. Our aggregation kinetics revealed that physiologically high concentrations of ATP facilitated the aggregation of both NM and FUS, which followed distinct aggregation mechanisms. FUS aggregation was facilitated by the high amount of ATP via phase separation, forming more liquid-like droplets that facilitated oligomerization and subsequent conversion to amyloid fibrils (15). Further, Mg^2+^ was found to be indispensable with ATP in modulating both of these aggregation kinetics. Our Raman data suggested that Mg^2+^ strengthened the electrostatic association between the anionic triphosphate moiety of ATP and positively charged amino acid residues in proteins such as lysine as suggested in previous reports. (14)(11)(48). Moreover, Mg^2+^ ions also possibly extended the interaction of ATP by acting as a bridge between negatively charged amino acids and triphosphates by minimizing their electrostatic repulsions (49). In order to gain more mechanistic insight into Mg^2+^-ATP dependent kinetic acceleration, we polymerized NM without ATP in the presence of high concentrations of MgCl_2_ to screen the repulsion between building blocks, which might promote the aggregation. However, a much higher Mg^2+^concentration was required to obtain a similar kinetic acceleration than ATP. Therefore, it possibly indicated the electrostatic crosslinking using lysine and multidentate triphosphate part of ATP that facilitated the critical nucleus formation, like in the case of the TauK18 aggregation (11). Additionally, the amphipathic nature of ATP allowed it to participate in different chemical interactions, similar to the industrial hydrotropes, utilizing its hydrophobic aromatic part to solubilize NM fibrils, as reported in a previous study (36). Interestingly, this dissolution of NM fibrils was found to be Mg^2+^-independent. Further, in agreement with the previous report, the extent of disaggregation was comparable, regardless of the ATP concentrations (5). In addition to these observations, we were also able to elucidate critical anti-prion-like attributes of molecular ATP, as represented in Figure 7. For example, ATP could not disaggregate NM-ATP amyloids polymerized in the presence of physiologically high concentrations of ATP, delimiting the number of autocatalytic amyloid seeds for the prion-like transmission. The trace ATP concentrations, reminiscent of oxidative stress or aging, gave rise to the seeding-incompetent NM amyloids that can impair the prion-like amplification cycles in recipient cells (50). Two principal factors of ATP governed these phenomena. Firstly, the extensive binding of ATP in the lysine-rich M-domain of NM reinforced an extensive crosslinking through ATP molecules, especially in high concentrations, that resulted in a protease-resistant core. This remarkable conformational compactness probably made these amyloids inaccessible to disaggregating factors such as free ATP or Hsp104 that target the M-domain (25)(51). Secondly, a remodeling in amyloid conformations in NM-ATP amyloids due to the binding of ATP during aggregation. Intriguingly, these nanoscale structural alterations in NM-ATP amyloids were dependent on the concentrations of ATP and were subject to the presence of Mg^2+^, as corroborated in our Raman and CD data. Our detailed analysis indicated that the seeding-competent NM-only and NM-ATP fibrils formed in the presence of 10 mM ATP had a higher ratio of β-sheet to the random coil as opposed to the other NM-ATP fibrils that showed a greater extent of a random coil. This reduction in the β-sheet content probably was the reason for their seeding inefficiency, except for the seeding-competent NM-ATP fibrils formed in the presence of 2.5 mM ATP (52)(53). In their case, the seeding inability due to the greater proportion of random coil was probably compensated by the extensive ATP binding in seeds, which can electrostatically attract the monomers to its growth-competent ends or surfaces. Moreover, the detailed profiling of ATP-disaggregated fibrils by AFM and immunoblots revealed that ATP did not promote fibril to oligomer formation, unlike the canonical disaggregase Hsp104. This pointed toward the fact that the disaggregation by ATP may have a limited effect on the transmission and pathology as oligomers are considered as the predominant particles for infection and toxicity in amyloid-associated diseases (54).

**Figure 7.**
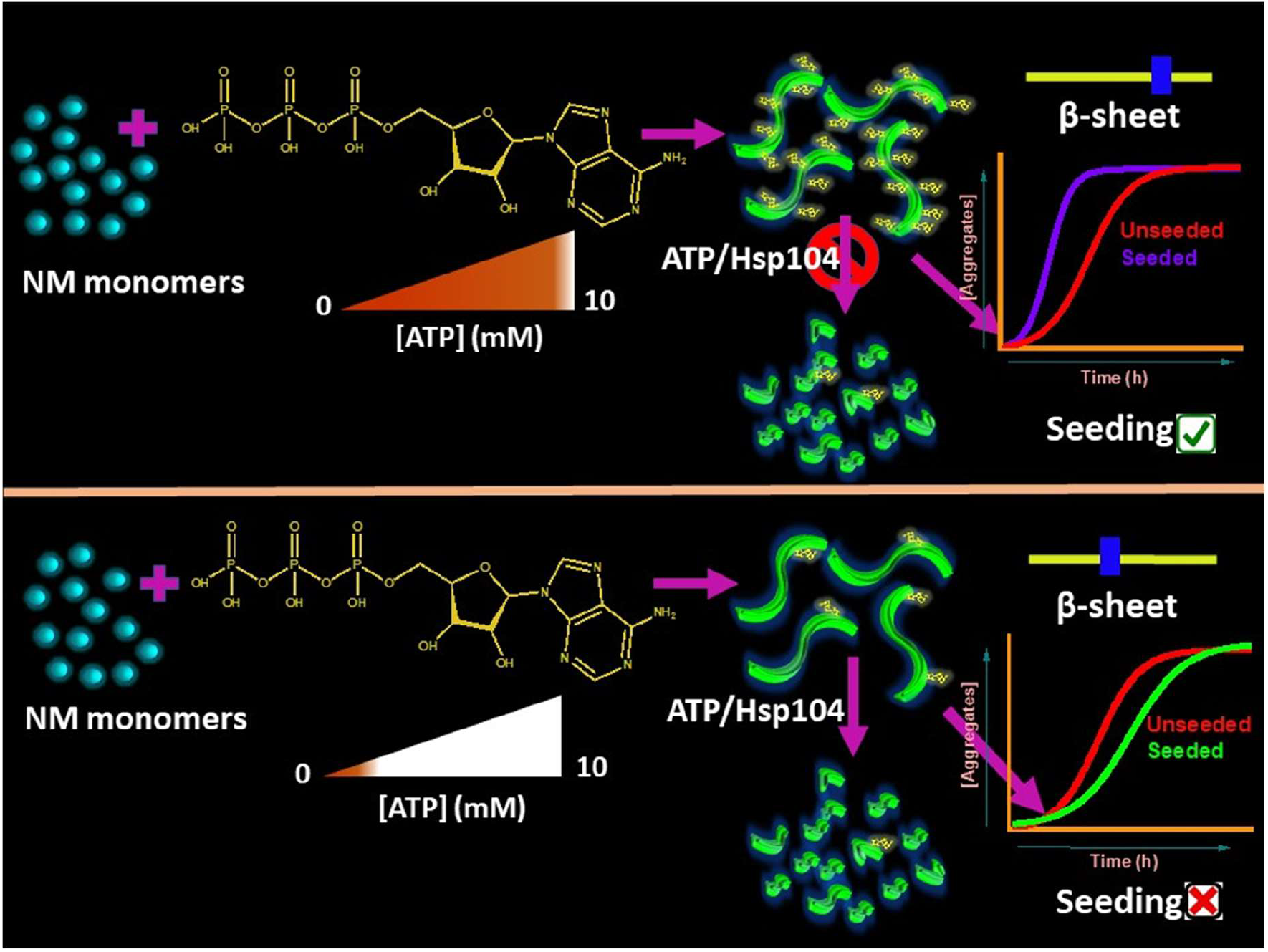
Proposed model showing anti-prion attributes of ATP molecules. ATP may restrict the prion-like amyloid transmission by creating ATP-bound, robust amyloids at physiologically high concentrations that cannot further fragment to generate seeds or by generating seeding-inefficient amyloids having reduced β-sheet content at trace concentrations.

In summary, our results provided key insights into ATP as a small chemical chaperone, that regulates the number and efficiency of seeds that may limit the spread of amyloids. Our findings on the modulation of amyloidogenesis covering orders of magnitude of the concentration range of ATP indicated some of the previously unexplored roles of this polyanion as an anti-propagation factor, in addition to its role in suppressing aggregation or dissolving preformed aggregates. Collectively, our study revealed a direct participation of molecular ATP to maintain proteostasis in addition to being the energy currency for chaperones. Considering generic nature of ATP-protein interactions, our results can have broad implications in functional non-Mendelian inheritance traits and transmissible neurodegenerative diseases associated with a diverse range of prions and prion-like domains.

## Materials and Methods

### Materials

Sodium phosphate dibasic dihydrate, magnesium chloride hexahydrate, Thioflavin-T (ThT), adenosine-5’-triphosphate disodium salt hydrate (ATP), dimethyl sulfoxide (DMSO), guanidinium hydrochloride (GdmCl), β-mercaptoethanol and HEPES [4-(2-hydroxyethyl)-1-piperazineethanesulfonic acid] were procured from Sigma (St. Louis, MO, USA). Urea and proteinase-K were bought from Amresco. Sodium dodecyl sulfate (SDS), imidazole, ammonium sulfate, lysozyme, potassium chloride, and ethylenediaminetetraacetic acid (EDTA) were purchased from HIMEDIA. Sodium hydroxide, potassium hydroxide, glycerol, methanol, nitrocellulose membrane, A11 (anti-amyloid oligomer antibody), OC (anti-amyloid fibril antibody), HRP-conjugated goat anti-rabbit antibody, and sodium chloride were bought from Merck. Antibiotics (ampicillin and chloramphenicol) and Isopropyl-β-thiogalactopyranoside (IPTG) were procured from Gold Biocom (USA). Q-sepharose and Ni-NTA columns were bought from GE Healthcare Lifesciences, USA. Fluorescein-5-maleimide was procured from Invitrogen. 96-well NUNC optical bottom plates were purchased from ThermoFisher Scientific (Waltham, Massachusetts, USA).

### Expression and Purification of NM

C-terminal hexa-histidine tagged recombinant Sup35NM protein was overexpressed in BL21 (DE3)/pLysS cells using IPTG, and then from harvested cells, proteins were extracted; the extracted proteins were subjected to first Ni-NTA purification by applying a gradient of imidazole and further from a Q-sepharose column using the gradient of sodium chloride. The detailed protocol was previously described by (25)(55). Also, for the single cysteine mutant of NM, the purification remains the same as mentioned above with the addition of β-mercaptoethanol.

### Site-directed mutagenesis for NM

Recombinant NM was cloned into the pET23a vector using appropriate primers. A single cysteine mutant was created using the site-directed mutagenesis kit, Quikchange (Stratagene). The following primers were used to create the mutation;

S150C Forward: GAAGCTTGTCTCCAGTTGCGGTATCAAGTTGG

S150C Reverse: GGCCAACTTGATACCGCAACTGGAGACAAGCTTC

DNA sequencing was performed to confirm the mutation.

### Expression and Purification of Hsp104

N-terminal His_6_-tag recombinant Hsp104 pPROEX-HTb-Hsp104 of *S. cerevisiae* was obtained from Addgene (61) and was overexpressed in BL21(DE3) RIPL *E. coli* cells using IPTG as inducer at 15 °C for 16 h. The detailed protocol for His_6_-Hsp104 purification and its subsequent His-tag removal is described by us (25)(56).

### Expression and Purification of FUS

MBP-*Tev-*FUS-*Tev-*His_6_ was overexpressed in *E. coli* BL21(DE3) RIPL strain, and the purification protocol was similar to that described earlier by us (57)(58).

### Amyloid aggregation reaction

Methanol-precipitated protein was dissolved and kept in a denaturation buffer (8 M Urea, 20 mM Tris-HCl, pH 7.4) at room temperature for at least 3 h. Denatured protein was passed through a 100-kDa filter first to remove any aggregates if present and then concentrated further using a 3-kDa filter for the amyloid aggregation reactions. The aggregation reactions were set up by incubating monomeric NM in assembly buffer (40 mM HEPES-KOH pH 7.4, 150 mM KCl, 1 mM DTT, 20 mM MgCl_2_, and 20 μM ThT) at room temperature and stirred at 80 rpm with the final protein concentration of 2.5 μM without or with different concentrations of ATP. ThT fluorescence was monitored at 450 nm excitation at room temperature, and emission was recorded at 480 nm. Also, NM aggregation reactions of 2.5 μM monomeric NM were set up in the presence of 10 mM ATP without MgCl_2_ in the assembly buffer under the same aggregation conditions. Alternatively, amyloid aggregation reactions of monomeric NM (2.5 μM) were carried out in 100 mM, 150 mM, and 250 mM MgCl_2_ (40 mM HEPES-KOH pH 7.4, 150 mM KCl, 1 mM DTT, and 20 μM ThT) buffer under the conditions as mentioned above.

### Seeded aggregation reaction

Amyloid seeds were generated by incubating monomeric NM (2.5 μM) in assembly buffer (40 mM HEPES-KOH pH 7.4, 150 mM KCl, 1 mM DTT, and 20 mM MgCl_2_) in the absence or presence of different concentrations of ATP at room temperature and stirring at 80 rpm. The resultant NM or NM-ATP amyloid seeds, after 6 h of aggregation reactions, were introduced to fresh aggregation reactions in assembly buffer (40 mM HEPES-KOH pH 7.4, 150 mM KCl, 1 mM DTT, 20 mM MgCl_2_, and 20 μM ThT) such that the seeds constitute 10% (w/w) of the total protein concentration (2.5 μM). The reactions were kept at room temperature and stirred at 80 rpm, and ThT fluorescence was recorded at specified time intervals. Alternatively, the NM-ATP amyloids generated in the presence of 2.5 μM ATP, as mentioned above, were buffer exchanged with buffer B (40 mM HEPES-KOH pH 7.4, 150 mM KCl, 1 mM DTT, and 250 mM MgCl_2_) and buffer A (40 mM HEPES-KOH pH 7.4, 150 mM KCl, 1 mM DTT, and 20 mM MgCl_2_), sequentially. These amyloids were introduced as seeds to fresh aggregation reactions in assembly buffer (40 mM HEPES-KOH pH 7.4, 150 mM KCl, 1 mM DTT, 20 mM MgCl_2_, and 20 μM ThT) as mentioned above, and ThT fluorescence was recorded with time. Also, amyloid seeds were generated by aggregating monomeric NM (2.5 μM) in assembly buffer (40 mM HEPES-KOH pH 7.4, 150 mM KCl, 1 mM DTT, and 20 mM MgCl_2_) at room temperature and stirring at 80 rpm, followed by sonication (Qsonica probe sonicator) at amplitude 5 for 2 pulses of 30 sec. These amyloid seeds were used in the seeded aggregation reaction, as mentioned above, by forming 10% (w/w) of the total protein concentration.

### Disaggregation of NM-only and NM-ATP fibrils

NM monomers (2.5 μM) were aggregated in assembly buffer (40 mM HEPES-KOH pH 7.4, 150 mM KCl, 1 mM DTT, 20 mM MgCl_2_, and 20 μM ThT) at room temperature and stirred at 80 rpm in the presence or absence of different concentrations of ATP. Different concentrations of ATP were introduced to the saturated aggregation reactions after 6 h of NM-only or NM-ATP aggregation, and a drop in ThT fluorescence was recorded at specific intervals. The ThT fluorescence intensities were normalized to the initial ThT fluorescence intensity before adding ATP. The percentage of disaggregation was calculated using [(Initial ThT fluorescence intensity -final ThT fluorescence intensity) / ThT fluorescence intensity] × 100% after 2 h. Alternatively, NM (2.5 μM) aggregation reaction was set up in the absence of MgCl_2_ in the buffer (40 mM HEPES-KOH pH 7.4, 150 mM KCl, 1 mM DTT, and 20 μM ThT) at room temperature and stirred at 80 rpm, and a drop in ThT fluorescence intensity with time on the addition of 10 mM ATP was recorded.

### Atomic force microscopy

Aggregation reactions of 2.5 µM of monomeric NM were set up in the assembly buffer (40 mM HEPES-KOH pH 7.4, 150 mM KCl, 20 mM MgCl_2_, 1 mM DTT) in the absence or presence of different concentrations of ATP, and samples were aliquoted after 6 h from the commencement of the reactions. Alternatively, NM-only fibrils (2.5 µM monomers) were disaggregated using different concentrations of ATP and also with 0.5 µM Hsp104 plus 5 mM ATP and ATP regeneration system (20 mM PEP and 15 µg/ml pyruvate kinase). Samples (10 µL) were deposited on a freshly cleaved mica and were then thoroughly washed with filtered water after 2 min of incubation with 200 µl of filtered water and later dried under a gentle stream of nitrogen. The AFM images were acquired on Innova atomic force microscopy (Bruker) using NanoDrive (v8.03) software. The images were processed and analyzed using WSxM 5.0 Develop 8.1 software (59), and height profiles were plotted using Origin.

### Sedimentation Assay

Monomeric NM (2.5 µM) was aggregated in the assembly buffer (40 mM HEPES-KOH pH 7.4, 150 mM KCl, 1 mM DTT, 20 mM MgCl_2_) at room temperature with stirring at 80 rpm. Multiple concentrations of ATP were added after 6 h from the commencement of the aggregation reaction and mixed thoroughly. Next, we pelleted amyloid species before and after the introduction of ATP at 16,400 rpm for 30 min. These pellets were resuspended in 8 M Urea (20 mM Tris-HCl pH 7.4) and incubated overnight to monomerize the amyloids. SDS-PAGE was performed, and band intensities were estimated using ImageJ software (60).

### The proteinase K digestion of fibrils

Monomeric NM (2.5 μM) was aggregated in the assembly buffer (40 mM HEPES-KOH pH 7.4, 150 mM KCl, 1 mM DTT, 20 mM MgCl_2_) without or with varying concentrations of ATP (2.5 nM, 2.5 μM, 2.5 mM, and 10 mM) for 6 h. The fibrils formed were pelleted by centrifugation at 16,400 rpm, 25 °C for 30 min, resuspended in the same buffer, and then incubated with proteinase K (PK) ([PK]: [Protein] 1:1000) at 37 °C for 30 min. Digestion reactions were stopped by adding SDS-loading dye and boiling them at 100 °C, followed by SDS-PAGE. Alternatively, the amyloid species retrieved from the pellets were resuspended in buffer B (150 mM KCl, 1mM DTT, 250 mM MgCl_2,_ and 40 mM HEPES-KOH pH 7.4) and were centrifuged at 16,400 rpm, 25 °C for 30 min. The retrieved pellets were resuspended again in the assembly buffer, followed by PK digestion and SDS-PAGE, as mentioned above.

### Dot blot assay

Aggregation reactions of monomeric NM (2.5 µM) were set up in the assembly buffer (40 mM HEPES-KOH pH 7.4, 150 mM KCl, 1 mM DTT, 20 mM MgCl_2_) at room temperature with stirring at 80 rpm. 2.5 nM or 10 mM ATP was introduced to the amyloids formed after 6 h from the commencement of the reaction. Alternatively, monomeric NM (2.5 µM) was aggregated in the assembly buffer (40 mM HEPES-KOH pH 7.4, 150 mM KCl, 1 mM DTT, 20 mM MgCl_2_) at room temperature with stirring at 80 rpm. Then after 6 h from the commencement of the reaction, Hsp104 (0.5 µM), ATP (5 mM), and ATP regeneration system (20 mM PEP and 15 µg/ml pyruvate kinase) were added. Two microlitres of samples were spotted on the nitrocellulose membrane before and after adding ATP or Hsp104. Next, for blocking, 3% BSA in PBST (0.05% Tween-20) was added and incubated for 1 h at room temperature, then blots were probed using a primary antibody (A11, 1:500; OC, 1:1000) overnight at 4 ºC. Next, the blots were washed six times with PBST and incubated with HRP-conjugated secondary antibody for 1 h at room temperature. The blots were thoroughly washed three times using PBST and subsequently developed using an ECL kit.

### Fluorescence labeling and confocal microscopy of FUS

Labeling reactions and microscopy were performed as described earlier (57). Confocal fluorescence imaging of FUS droplets and aggregates was performed on ZEISS LSM 980 Elyra 7 super-resolution microscope using a 63x oil-immersion objective (Numerical aperture 1.4). For visualizing droplets of FUS, 200 nM of Alexa488-labeled protein was added with 10 µM unlabeled protein, and 10 μL of the freshly phase-separated sample was placed on a coverslip. ThT-bound aggregates were directly visualized under the same settings used for droplets. ImageJ (NIH, Bethesda, USA) software was then used for processing and image analyses (57)(60).

### Phase separation and aggregation assays of FUS

Droplet formation of FUS were initiated TEV protease cleavage in a 1:10 molar ratio (TEV: protein) in 20 mM sodium phosphate, pH 7.4 buffer at room temperature. Turbidity of phase-separated samples was done by recording the absorbance at 350 nm using 96-well NUNC optical bottom plates on a Multiskan Go (Thermo Scientific) plate reader. For aggregation reactions, FUS phase separation reactions (10 µM of FUS with 20 µM ThT) were stirred at 600 rpm after 15 min of TEV addition. Aggregation kinetics was then monitored as a function of ThT fluorescence at 480 nm with time.

### Fluorescence recovery after photobleaching (FRAP) measurements

FRAP experiments for droplets with and without ATP were performed on ZEISS LSM 980 Elyra 7 super-resolution microscope using a 63x oil-immersion objective (Numerical aperture 1.4). Alexa488-labeled protein (200 nM) was mixed with 10 µM of unlabeled protein, and droplet formation was induced by TEV. ZEN Pro 2011 (ZEISS) software (provided with the instrument) was used to record the recovery of the bleached region, which was then normalized and plotted after background correction using Origin.

### Raman spectroscopy

NM monomers were aggregated in assembly buffer (40 mM HEPES-KOH pH 7.4, 150 mM KCl, 1 mM DTT, 20 mM MgCl_2_) without or with several concentrations of ATP (2.5 nM, 2.5 μM, 2.5 mM, and 10 mM) at room temperature with stirring at 80 rpm for 6 h. Alternatively, monomeric Sup35 NM (2.5 µM) were aggregated in the buffer (40 mM HEPES-KOH pH 7.4, 150 mM KCl, 1 mM DTT) with and without 20 mM MgCl_2_ in the presence of 10 mM ATP at room temperature with stirring at 80 rpm for 6 h. Next, the amyloid species were pelleted after 6 h at 16,400 rpm for 30 min and resuspended in 5 mM Na_2_PO_4_, 150 mM NaCl pH 7.4 buffer. Three μL of the pelleted sample was deposited onto a glass slide covered with aluminum foil and dried under a gentle stream of nitrogen gas. This step was repeated thrice. The sample was focused using a 50x objective lens (Nikon, Japan). A NIR laser, 785 nm with an exposure time of 10 sec and 250 mW laser power, was used to excite the samples. All the spectra were averaged over 20 scans. Rayleigh scattered light was removed using an edge filter, while the Raman scattered light was dispersed using a 1200 lines/mm diffraction grating and was further detected using an air-cooled CCD detector. Wire 3.4 software provided with the instrument was used for data acquisition, and the recorded Raman spectra were baseline corrected using the cubic spline interpolation method and smoothened in the same software. Spectra were finally plotted using Origin.

### Circular dichroism (CD) measurements

Monomeric NM (2.5 µM) was aggregated in assembly buffer (40 mM HEPES-KOH pH 7.4, 150 mM KCl, 1 mM DTT, 20 mM MgCl_2_) without or with several concentrations of ATP (2.5 nM, 2.5 μM, 2.5 mM, and 10 mM) at room temperature with stirring at 80 rpm for 6 h. The amyloid species were pelleted after 6 h at 16,400 rpm for 30 min and resuspended in 10 mM Na_2_PO_4_ (pH 7.4) buffer. The final concentration of NM was ∼8 µM, after which far-UV CD spectra were recorded on a Chirascan CD spectrometer (Applied Photophysics, UK) at room temperature. All the spectra were collected and recorded in the scan range 195-260 nm with a 1 nm step size and a quartz cuvette of 1 mm path length. The spectra were averaged over 5-10 scans, and the buffer signal was subtracted. All the spectra were smoothened using the ProData software provided with the Chirascan CD Spectrometer. Finally, the mean residue ellipticity [θ] was calculated, and plots were generated using Origin software.

### Fluorescence labeling of NM

Cysteine mutant of NM was labeled in the denaturation buffer 8 M Urea (20 mM Tris-HCl pH 7.4). A 30 mM stock of Fluorescin-5-maleimide was prepared in DMSO and was mixed in a ratio of 1:10 with the NM mutant. The reaction mixtures were kept in the dark for 2-3 h at room temperature on rotor spin. Next, to remove excess dye, the labeled protein was then buffer exchanged in a 3-kDa filter using 8 M Urea (20 mM Tris-HCl pH 7.4). The concentration of labeled protein was estimated using ɛ_494_ = 68000 M^−1^cm^−1^, and the labeling efficiency was ˃90% for the mutant protein.

### Steady-state fluorescence measurements

Steady-state fluorescence measurements of a single cysteine mutant of NM (S150C) were carried out using a FluoroMax-4 spectrofluorometer (Horiba Jobin Yvon, NJ) at room temperature quartz cuvette of 1 mm path length. For recording the fluorescence spectra, the mutants were excited at 485 nm, where the excitation and emission slits were 1.75 and 5 nm, respectively. Concomitantly, steady-state anisotropy measurements were performed by setting the excitation wavelength at 485 nm and emission wavelength at ∼518 nm with excitation and emission slits as 2 and 8 nm, respectively.

### Statistical analysis

All the experiments were performed three times, and the data are represented as mean ± SD indicated by scattered data points from independent experimental replicates. The statistical significance analysis was performed using one-way ANOVA tests, and the *p*-values were indicated in the figure legends. All the data analysis, data fitting (adjusted R^2^ > 0.95), and data plotting were performed with the help of Origin.

## Data Availability

All data described are contained within this paper and Supporting Information.

## Supporting Information

This article contains Supporting Information.

## Acknowledgments

We thank IISER Mohali, Department of Biotechnology (grant to S. Mukhopadhyay), Department of Science and Technology (INSPIRE fellowship to S. Mahapatra) for financial support, Dr. Dominic Narang (Mukhopadhyay lab) for the plasmid used in this work, Prof. Dorothee Dormann (Ludwig-Maximilians University and Institute für Molekulare Biology, Mainz) for her kind gift of FUS plasmid, Ms. Lisha Arora for drawing the structures, and the members of the Mukhopadhyay lab for critically reading this manuscript.

## Author Contributions

S. Mahapatra conceived the research and designed the study. S. Mahapatra and A.S., N.P., A.J., A.A., and A.W. performed the experiments and analyzed the data. S. Mahapatra, A.S., and N.P. wrote the first draft. S. Mukhopadhyay edited the manuscript, supervised the work, and obtained funding. All authors discussed the results and commented on the manuscript.

