## Supplementary Information for "ATP modulates self-perpetuating conformational conversion generating structurally distinct yeast prion amyloids that limit autocatalytic amplification"

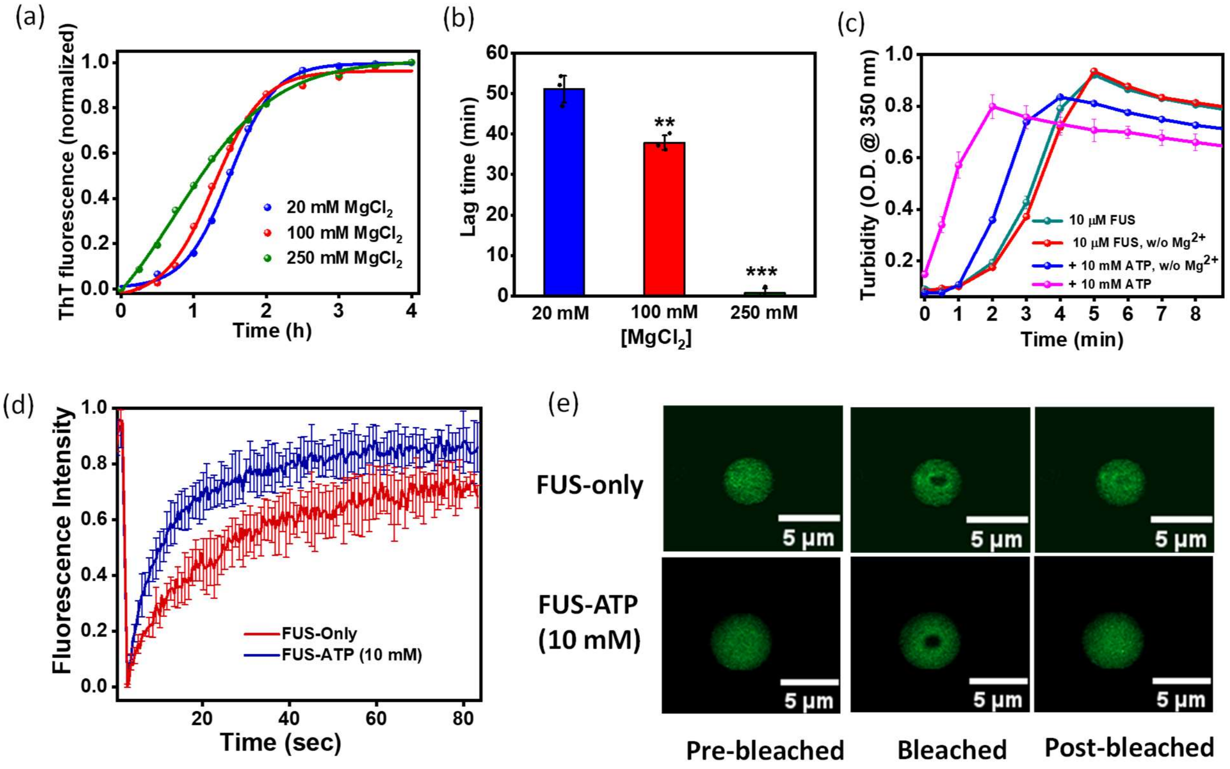


**Figure S1.** (a) Representative normalized ThT fluorescence kinetics of rotated (80 rpm) NM (2.5 μM) aggregation in the presence of 20 mM, 100 mM, and 250 mM $MgCl_2$ at room temperature. (b) Lag times of rotated (80 rpm) aggregation of NM monomers (2.5 μM) in the presence of 20 mM, 100 mM, and 250 mM $MgCl_2$ at room temperature. Standard deviations were calculated from three independent replicates (n = 3), $P < 0.01$, $P < 0.001$ for lag times in presence of 100 mM and 250 mM $MgCl_2$, respectively, compared to the lag time in the presence of 20 mM $MgCl_2$. (c) Liquid-liquid phase separation of FUS (10 μM) in the absence of ATP. Phase separation of FUS (10 μM) without $MgCl_2$.Also, the liquid-liquid phase separation of FUS (10 μM) in the presence of 10 mM ATP in the absence or presence of $MgCl_2$ was monitored by the turbidity assay (at 350 nm). Standard deviations were calculated from three replicates (n = 3). (d) FRAP kinetics of FUS (10 μM) droplets formed in the absence or presence of 10 mM ATP after 15 min. The solid lines represent the fitted curves. (e) The fluorescence images of FUS or FUS-ATP (10 mM) droplets during FRAP measurements are shown.

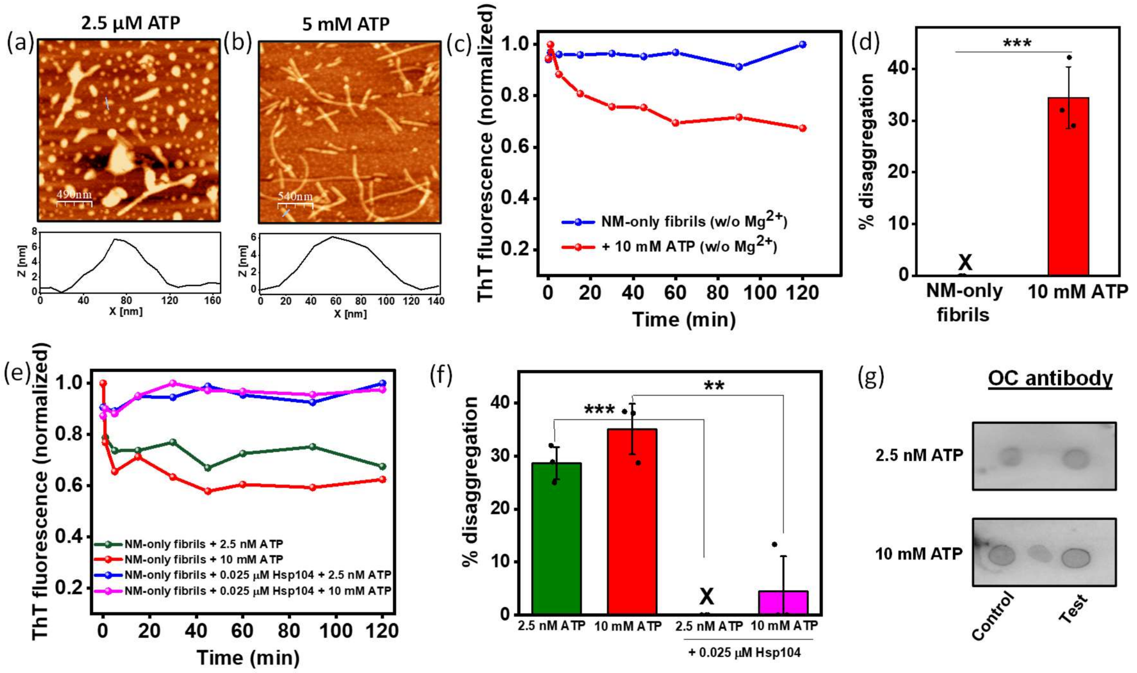


**Figure S2.** (a,b) AFM images showing disaggregation of amyloids formed from rotated (80 rpm) aggregation reaction of NM (2.5 μM) at room temperature by (a) 2.5 μM ATP and (b) 5 mM ATP with a height of ~6 nm. (c) Representative disaggregation kinetics of NM-only fibrils (2.5 μM monomers) without or with 10 mM ATP in the absence of $MgCl_2$ at room temperature and 80 rpm, showing the percentage of disaggregation in Figure S2d. Standard deviations were calculated from three independent replicates (n = 3), $P$ < 0.001 for disaggregation by 10 mM ATP compared to NM-only fibrils. (e) Representative disaggregation kinetics of NM-only fibrils (2.5 μM monomers) by ATP in the absence or presence of Hsp104 (0.025 μM) at room temperature and 80 rpm. (f) Percentage of disaggregation of NM-only fibrils (2.5 μM monomer) formed at room temperature and 80 rpm by ATP in the absence or presence of Hsp104 (0.025 μM). Standard deviations were calculated from three independent replicates (n = 3), $P$ < 0.001 for disaggregation by 2.5 nM ATP in the presence of Hsp104 compared to in the absence of Hsp104; $P$ < 0.01 for disaggregation by 10 mM ATP in the presence of Hsp104 compared to in the absence of Hsp104. (g) NM-only fibrils (2.5 μM monomer) were spotted before and after disaggregation by 2.5 nM and 10 mM ATP on the nitrocellulose membrane and were probed using the OC antibody.

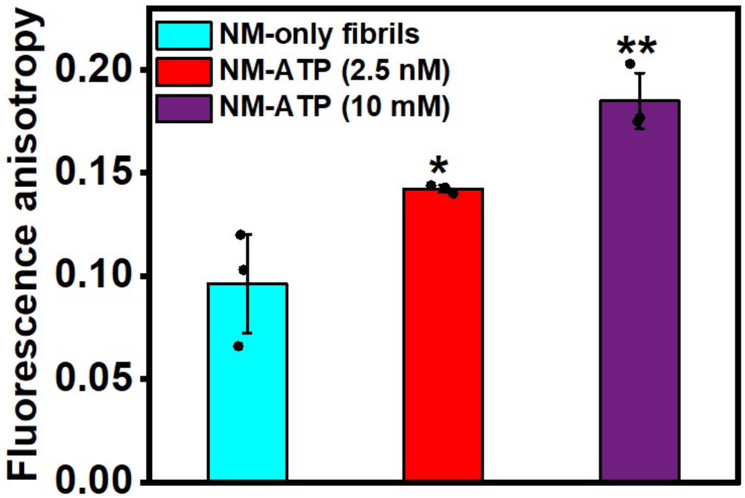


**Figure S3.** Steady-state fluorescence anisotropy of fluorescein-5-maleimide-Cys150-NM amyloid fibrils formed in the absence or presence of different concentrations of ATP. Standard deviations were calculated from three independent replicates (n = 3), $P < 0.05$ in the presence of 2.5 nM and $P < 0.01$ in the presence of 10 mM ATP compared to NM-only fibrils.

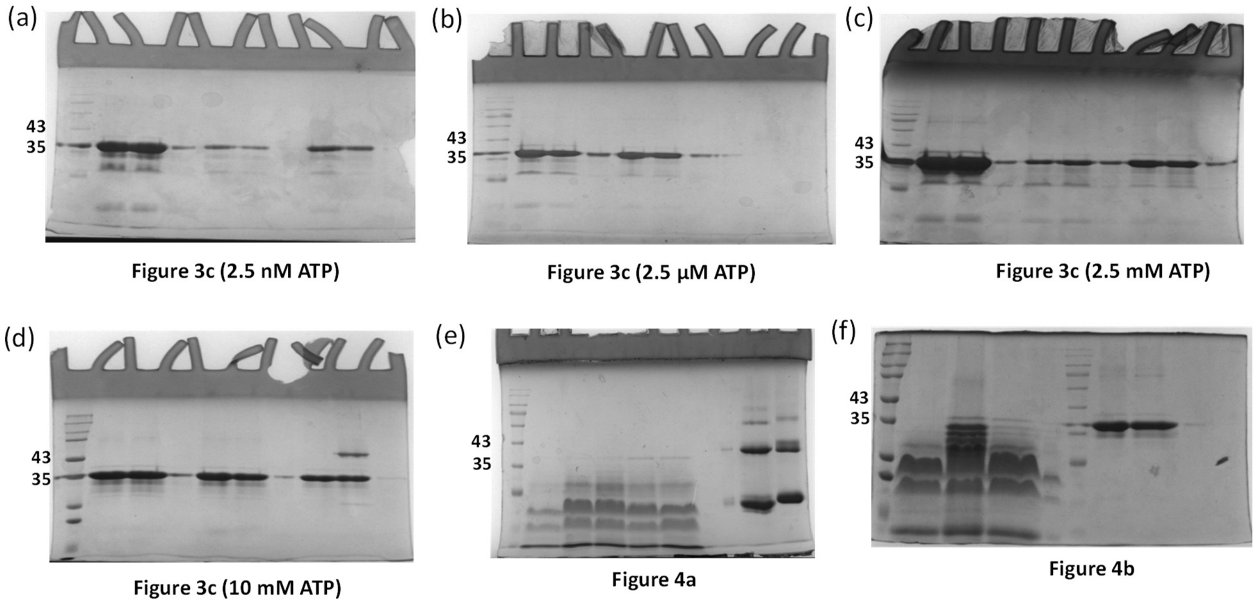


**Figure S4.** Full uncropped gels shown in Figures 3 and 4.
